# Single cell Transcriptome and T cell Repertoire Mapping of the Mechanistic Signatures and T cell Trajectories Contributing to Vascular and Dermal Manifestations of Behcet’s Disease

**DOI:** 10.1101/2022.03.22.485251

**Authors:** Ling Chang, Zihan Zheng, Qinghua Zou, Bing Zhong, Chengshun Chen, Xian Cheng, Qingshan Ni, Tiantian Che, Zhihua Zhao, Chunhao Cao, Yiwen Zhou, Xiangyu Tang, Zhifang Zhai, Jing Zhao, Junying Zhang, Liting Wang, Ying Wan, Guangxing Chen, Jingyi Li, Liyun Zou, Yuzhang Wu

**Author notes:** These authors contributed equally to the manuscript (Ling Chang, Zihan Zheng).

## Abstract

Behcet’s disease (BD) is a form of vasculitis characterized by complex multi-organ manifestations that may frequently recur and induce major tissue damage. Although genetic association studies have identified a number of risk factors, the etiology of BD and its tissue manifestations remains unknown, and the landscape of immune responses in BD is opaque, particularly in terms of inflammatory recurrence. In this study, we mapped the transcriptomes of the immune cell compartment in BD at single-cell resolution, sampling both circulation and affected skin in order to chart the immune interplay driving pathogenesis. Through comprehensive expression and communication analysis of the twenty major cell types identified, we observe striking mechanistic differences in immune response between BD skin lesions and peripheral circulation involving TNF signaling and T cell migration. Through integrated TCR sequencing, we further discover a pattern of clonal sharing between circulating and skin CD8+T cell populations along a trajectory defined by the acquisition of tissue-residential properties. In addition, we also identify a population of expanded CD4+ Tregs with the propensity to produce IL-32. Instead of suppressing effector T cell proliferation and function, IL-32 triggers increased expression of CD97, and may thus encourage prolonged local T cell activity in the skin. Collectively, our data serve to advance understandings of contributions of varying immune cell types to BD pathogenesis in the vasculature and skin, as well as the lifecycle patterns of T cells clones in this context. These data may also assist in further investigations of the mechanisms contributing to Treg dysfunction in systemic autoimmunity, while generating a conceptual model of T cell function contributing to BD recurrence.

## Introduction

Behcet’s disease (BD) is a recurrent form of vasculitis characterized by complex, multi-organ, manifestations involving mucosal membranes, skin, eyes, and other tissues^1^. These inflammatory manifestations often run a course of remission and relapse, and may cause significant tissue damage and negatively impact patient quality of life^2^. While the precise etiology of BD remains unclear, aberrant immune responses has long been appreciated to be a dominant contributor, with early genetic linkage studies demonstrated an association between class I major histocompatibility (MHC) complex molecules and BD, particularly *HLA-B*51*^3,4^. Larger-scale genome-wide association studies have confirmed this association, and have further identified a number of other risk alleles impacting genes responsible for modulating immune response, such as *IL10, IL23R-IL12RB2*, and *IFNGR1*^5,6,7^. Functional studies of immune cells isolated from BD patients have also reported altered behavior across multiple populations, including increased T cell oligoclonal expansion^8^ and cytokine production^9^, increased memory B cells of certain isotypes^10,11^, and M1-skewing of macrophages^12^, among many others. However, many of these studies have focused on changes in peripheral circulation, and are limited in scope to pre-defined cell populations, while a fuller census of the cell types involved is currently lacking. Furthermore, it is unclear how these different cell types interact with each other to drive vasculitis, and whether these patterns of behavior are also responsible for mediating the inflammatory organ manifestations seen in BD.

While simultaneous assessments across differing immune cell populations has traditionally been arduous due to the throughput limitations, advances in library construction methodologies in recent years have made it increasingly possible to concurrently profile transcriptomes of distinct cell types at single-cell resolution and census cell types in a systematic manner^13^. Further technological advances have been made it possible to recover of specific T cell receptor (TCR) and B cell receptor (BCR) sequences in single cells to simultaneously classify the adaptive immune repertoire in complex samples^14^. While these technologies have already been applied to classify immune cell type heterogeneity and T cell clonal behavior in multiple contexts, including cancer^15^, and infection^16^, these tools have yet to be fully applied in the context of autoimmunity and rheumatologic disorders such as BD^17^. In this manuscript, we utilize scRNAseq and scTCRseq to map for the first time the immune cell types present in peripheral circulation and inflamed skin of patients with BD. We perform detailed functional analysis of the molecular pathways, cell types, and communication patterns showing differential regulation between skin lesions and circulating immune cells in BD, both compared internally and as compared to similar cell types identified in healthy controls. We also develop a novel computational approach to trace the behavior of CD4+ and CD8+ T cell clones across circulation and skin in BD. The unified analyses presented here may serve as a valuable resource for understanding the behavior of specific immune cell types that contribute to BD pathogenesis.

## Materials and Methods

### Functional analysis of single-cell data

Functional analyses of pathway enrichment were performed using the GSEA method^18^ using the Reactome knowledgebase^19^. Since droplet-based scRNAseq data tends to show sparse expression in individual cells, each cluster was sampled 5 times to generate bulk profiles of 100 cells each for more accurate comparisons. Functional communication analyses were performed using the CellChat package^20^ in R using the built-in human receptor-ligand reference database. Trajectory inference was performed using the Slingshot package^21^ in R run on the first two dimensions of UMAP embedding. Identification of genes displaying significant changes in expression over the course of pseudotime progression was performed using the TIPS package^22^ in R. Network visualization of the expression correlation between progression-associated genes was obtained through Fruchterman-Reingold force-directed expansion in Gephi. Other visualizations were drawn using the ggplot2^23^, UpSetR^24^, and pheatmap^25^ packages in R.

### Integrated analysis of scTCRseq data

To integrate TCR sequences with transcriptome data, we first filtered out cells with multiple distinct TCRβ and/or >2 TCRα chains as potential doublets, and matched the remaining TCR sequences to their originating cell based on cell barcode. Clonal homeostasis analysis within specific T cell subsets was performed by tabulating the number of cells corresponding to a single TCRβ clonotype in each population. Morista-Horn similarity between cell types was computed based on cell counts of each clonotype using the abdiv package^26^ in R. TCR clustering was performed using the GLIPH2 algorithm^27^ with full substitution matrixes.

### Flow Cytometry

PBMCs were isolated from peripheral blood of healthy donors by density gradient separation (Lymphoprep, Stem Cell Technologies) and cryopreserved in 90% FBS (Gibco) and 10%DMSO (Sigma-Aldrich). Prior to experimental manipulation, PBMCs were recovered and rested overnight. Single-cell suspensions of rested PBMCs were aliquoted into plates pre-coated with anti-CD3 (clone : UCHT1, Biolegend) and anti-CD28 (Clone:CD28.2, Biolegend) antibodies according to manufacturer’s protocol at a density of 1X10^6^/mL to induce TCR activation. 2000ng/mL recombinant human (rh) IL32γ (4690-IL-025/CF, R&D Systems) or solvent control was given at the same time, and cells were incubated at 37°C in 5% CO_2_ and 100% humidity in RPMI-1640 supplemented with 10% FBS and 100ug/ml penicillin/streptopmycin (Gibco).

For intracellular cytokine production experiments, cells were maintained for 3 days, and further stimulated with a cocktail of (1 μg/mL Iomomycin; 50 ng/mL phorbol-12-myristate 13-acetate (PMA); 3 μg/mL Brefeldin A (BFA)) for 4 hours priors to collection. Cells were then harvested and stained with a amine-reactive dye (NIR, Invitrogen) and surface antibodies against CD3, CD4, and CD8 (BD Bioscience). Intracellular staining (BDB554714, BD) was performed according to the manufacturer’s protocol using antibodies against TNF, IL2, GZMB (BD Bioscience), and IFNγ (Invitrogen). For cell proliferation experiments, cells were labeled with Celltrace (1922850, Invitrogen) according to the manufacturer’s protocol. Cells were then incubated for 6 days prior to collection, and stained with antibodies against CD3, CD4, CD8, CD97 (BD Biosciences), PD1, CXCR6, and CD57 (Biolegend).

### Human Treg Differentiation

CD4+ naive T cells were isolated from healthy donors’ peripheral blood by CD4 MicroBeads according to the manufacturer’s protocol (Miltenyi). A non-differentiated control group (Th0) was treated with anti-CD28 Ab and rhIL2 (Biolegend) alone, while Tregs were differentiated through treatment with anti-CD28 Ab, rhIL2 and rhTGFβ (LS-G4063, R&D Systems). Th0 and Treg groups were also exposed to rhIL32γ to assess the impact of the cytokine on differentiation efficacy. Cells were harvested at 3 days and stained with surface antibodies against CD4 and CD25 (BD Bioscience), processed using a transcription factor buffer kit (Biolegend), and subsequently stained with antibody against FOXP3 (Biolegend).

## Results

### Immune cell atlas of Behcet’s disease

In order to more completely characterize the immune response underlying Behcet’s disease, we first performed paired scRNAseq and scTCRseq on peripheral blood of four patients and affected skin samples from two patients (Fig1A). Stringent filtering based on quality control metrics yielded a total of 39,820 high-quality single-cell transcriptome profiles (FigS1A). Following dimension reduction and clustering, 20 major cell populations were identified and annotated based on their expression profiles (Fig1B, TableS1). These population-characteristic genes were both highly expressed and largely cluster-unique (Fig1C, TableS2).

**Figure 1.**
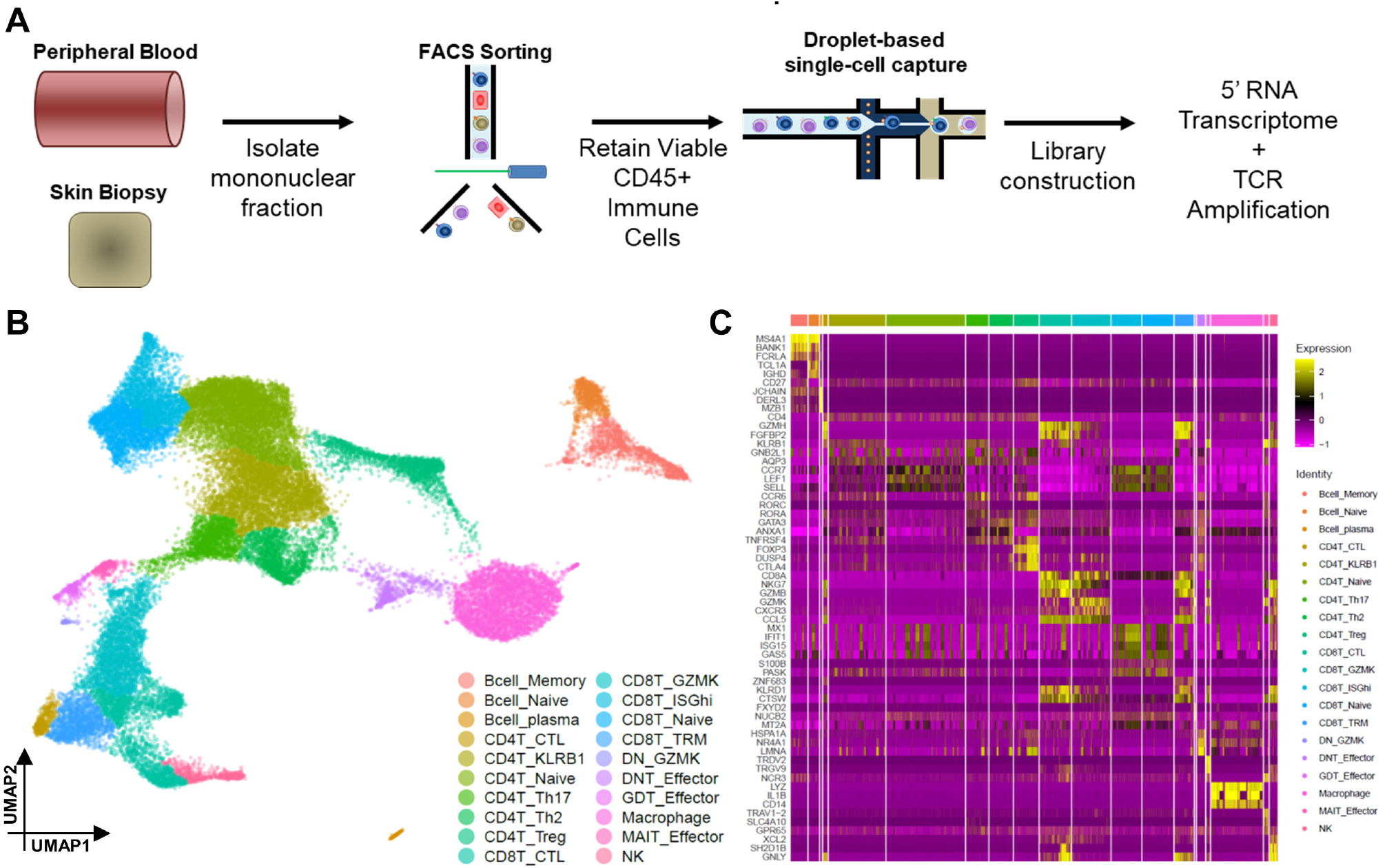
Single cell RNAseq atlas of immune cell populations in peripheral blood and skin lesions of Behcet’s Disease patients. A) Schematic overview of the experimental design for single-cell sequencing. Mononuclear cells isolated from peripheral blood and skin biopsies of BD patients with active disease were sorted through FACS, and CD45+ immune cells were retained for droplet-based single-cell capture and subsequent library construction. In order to simultaneously capture TCR sequences and transcriptome information, 5’ sequencing was performed. B) UMAP visualization of the 20 cell populations identified following dimension reduction and clustering. C) Heatmap visualization of select representative markers of each cell population. While these markers were selected based on expression differential against all other cell types, some markers may be expressed in several relatively similar populations.

Many of these populations have been previously described to have altered counts and/or function in BD. For instance, three populations of B cells were identified, corresponding to naïve, memory, and plasma phenotypes. Consistent with previous reports, we observed a high ratio of memory cells to naïve cells in both the blood and skin of BD patients (FigS2), with a substantial proportion of these memory cells being IgA-secreting clones. At the same time, we were also able to capture rarer cell populations, including γδT, which have been suggested to be functionally restrained in BD^28^, as well as natural killer cells, whose potential contribution remains to be explored^29^. We also observed significant numbers of rare T cell types, including mucosal-associated invariant T cells reactive to antigen presented by MR1 and which have been suggested to contribute to inflammatory pathogenesis in other forms of vasculitis^30,31^, as well as double-negative αβT cells expressing neither CD4 nor CD8, which have been previously reported to be increased in BD^32^. By capturing the transcriptome profiles of significant numbers of these rare cell types, together with larger macrophage and conventional T cell populations in both skin and circulation, we are thus able to create an unbiased census of the cell types potentially involved in both vasculitis and skin manifestations. This survey can consequently serve as the basis for the further functional and lineage analyses into BD pathogenesis.

### Genetically-associated markers of Behcet’s disease are abundantly expressed

A substantial number of studies have been performed to identify risk alleles and genes associated with susceptibility to BD across multiple populations, but the expression distribution of many of these identified markers across immune cell populations has been hitherto unclear. At the same time, other studies have identified specific functional aberrations in specific populations in BD, but have been unable to address their relation to BD risk alleles. Taking advantage of the resolution offered by scRNAseq, we mapped the expression levels of genetically-associated risk genes across the cell types identified in our dataset (Fig2A, FigS3) to synthesize these two lines of knowledge. Interestingly, we observed that many markers demonstrated focused expression, such as *CCR1, TLR4, NOD2*, and *MEFV*, which were almost exclusively found in macrophages of both skin and peripheral circulation (Fig2B). Similarly, the transcription factor *FOXP3* was exclusively expressed in CD4+ Tregs, and *FCRL3* was only found in B cells and some CD4+T populations. At the same time, other genes, including *TNFAIP3, HLA-B, UBAC2, STAT4*, and *GIMAP4* were highly expressed and detectable across a large majority of the immune populations analyzed. As such, these patterns suggest that both defects in specific populations and more general mutations can contribute to BD pathogenesis.

**Figure 2.**
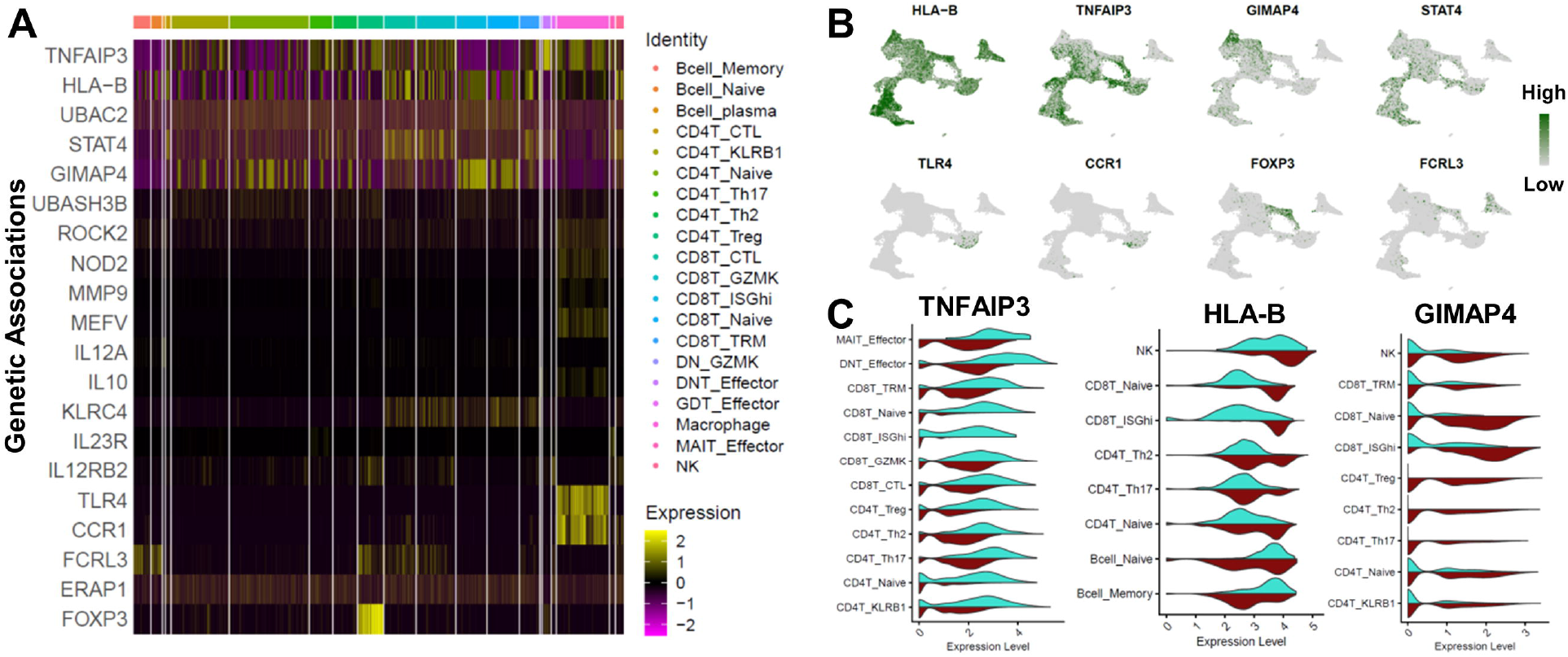
Genetically-associated markers of Behcet’s Disease are abundantly expressed in both peripheral and skin populations. A) Heatmap of the expression of 20 markers reported in genetic-association studies to include alleles associated with BD risk, and which were found to be significantly expressed in at least one of the immune cell populations observed. B) Expression profiles of eight genetically-associated markers projected in UMAP space. C) Violin plots comparing the expression of three markers (*TNFAIP3, HLA-B*, and *GIMAP4*) among skin and immune cell populations. Cell types not found to express these markers in either tissue were excluded. Turquoise halves correspond to marker expression in skin, while garnet halves correspond to expression in circulation.

Interestingly, some of these broadly expressed markers also showed differential expression between peripheral blood and skin populations. In particular, expression of *TNFAIP3* (encoding for the key signaling adaptor A20) was elevated in skin T cell populations compared to their peripheral blood counterparts, while expression of *GIMAP4* was significantly lower. *TNFAIP3* has been demonstrated to serve as a negative regulator of inflammation by being induced by TNF, and subsequently restraining, TNF-driven apoptosis via NF-kB. Intriguingly, mutations in *TNFAIP3* associated with BD have been reported to involve a loss-of-function in its suppressive capability. These indications thus raise the possibility that the increase of *TNFAIP3* in skin populations may be the result of active TNF stimuli, but that the expressed protein may not be sufficiently potent to suppress TNF downstream effects. At the same time, *GIMAP4* has previously been reported to regulate the secretion of the IFNγ^33^ and decrease during differentiation towards Th2 cells^34^, suggesting that downregulation of this factor may be involved in enabling differentiation of CD4+T cells towards non-Th1 fates observed in the skin during BD.

### Differential characteristics between skin and peripheral immune cells in Behcet’s Disease

Having already observed some significant differences between skin and peripheral immune cell populations within the genetically-associated markers analyzed above, we then sought to comprehensively catalogue these changes over the entire transcriptome. Labeling of individual cells in the UMAP space based on tissue origin revealed that the relative percentages of many populations varied significantly (Fig3A). Of note, we observed that naïve CD8+T cells and ISGhi CD8+T cells were very rarely observed in the skin, while CD4+T Tregs and DN effector T cells were prominently enriched therein (Fig3B). A significantly lower percentage of macrophages were also found in the skin, while relative proportions of B and NK populations were relatively similar.

**Figure 3.**
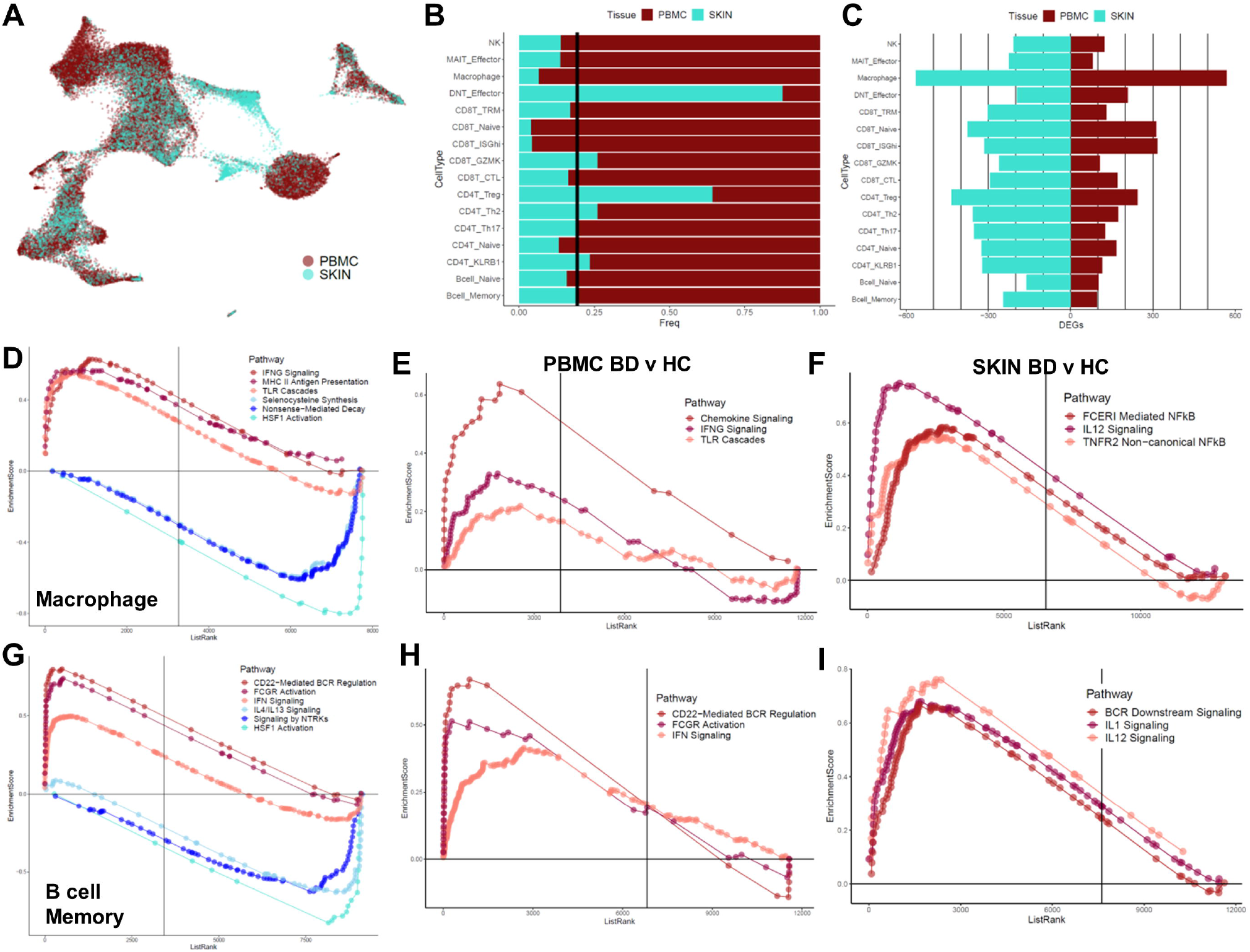
Differential expression characteristics between skin and peripheral immune cells in Behcet’s Disease. A) Visualization of the distribution of skin (turquoise) and circulating (garnet) cell populations in UMAP space. While cells from both tissues are largely interspersed, some populations notably appear to be overrepresented by one tissue. B) Barplot of the proportional representation of skin and circulating cells in each cell type. Due to differences in absolute cell count, a reference line is depicted to demarcate the point of equal representation based on relative likelihood. C) Pyramid plot depicting the numbers of genes showing significant differential expression between in skin and circulation within each cell type. D-F) Gene set enrichment plots showing functional pathways with differential expression in macrophages in circulation versus skin of BD patients (D), enriched in circulating macrophages in BD patients compared to healthy controls (E), and enriched in skin macrophages in BD patients compared to healthy controls (F). G-I) Gene set enrichment plots as in (D-F), comparing functional changes in memory B cells.

Interestingly, differential expression analysis demonstrated that individual cell populations also tended to show substantial transcriptome differences between skin and peripheral blood (Fig3C). While a few of these differences were shared across multiple populations at statistically significant levels (FigS4), the majority were constrained to one or two, indicating that these differences were unlikely to result from systemic biases in sequencing. Functional analyses of these differences in macrophages, the population with the greatest apparent variation, demonstrated that circulating macrophages showed elevated interferon signaling and toll-like receptor (TLR) signaling, as well as increased expression of MHC Class II antigen presentation machinery (Fig3D). Similarly, circulating memory B cells showed higher interferon signaling and FCGR activation compared to their counterparts in the skin (Fig3G). These results, coupled with the higher proportions of circulating ISGhi CD8+T cells observed, suggested that interferon activity may figure prominently in driving the underlying vasculitis phenotype contributing to BD, but may be less critical to its dermal pathogenesis.

To further evaluate this hypothesis, we then performed enrichment comparisons against macrophages and B cells derived from healthy controls in a publicly available dataset^35^. Consistent with our expectations, circulating macrophages from BD patients had increased enrichment for pathways associated with interferon signaling, TLR signaling, as well as chemotaxis (Fig3E). Similarly, circulating memory B cells from BD patients displayed enriched interferon signaling, FCGR activation, and BCR regulation compared to those from healthy controls (Fig3H). Collectively, these results support our observation that inferno activity may intimately affect the functions of multiple circulating immune cell populations in BD.

At the same time, macrophages derived from the skin of BD patients showed elevated activity in NF-kB as mediated through antibody receptor signaling, as well as TNFR2, together with increased IL-12 signaling (Fig3F). Furthermore, skin memory B cells showed prominent alterations in IL-1, IL-12, and BCR downstream signaling (Fig3H-I). These results indicate that circulating and skin macrophage and memory B cell populations are functionally distinct from their counterparts in healthy skin and circulation, and furthermore distinct in different ways. Furthermore, the skin environment appears to significantly reshape these inflammatory signatures seen in the periphery towards a local response profile, with a potential sensitivity to NF-kB activation.

In order to further understand the mechanisms contributing to local education of immune cell populations in the skin of BD patients, we then categorized the differential communication patterns between immune populations in the skin and peripheral circulation based on well-annotated receptor-ligand interactions. UMAP classification of these interactions demonstrated showed that many of these interactions were shared in both tissues on a general level (Fig4A, FigS5-6). In agreement with our observations above, ISGhi CD8+T cells appear to play a significant role in mediating immune cell crosstalk in peripheral circulation where they appear in large numbers, but not in skin tissue where they are scarce (Fig4B). On the other hand, a number of signaling interactions were enriched in skin compared to peripheral circulation. For instance, we could observe highly elevated TNF communication among skin immune populations (Fig4C), consistent with the increase in TNFAIP3 expression in the skin we observed above. TNF has long been suspected to play a role in the pathogenesis of BD, and treatment using anti-TNF antibodies has been demonstrated to have considerable efficacy in BD cases with skin and mucosal involvement^36^. Signaling via *ADGRE5* (encoding for CD97), a surface molecule known to be involved in immune cell migration^37,38^, was also increased among skin immune populations (Fig4D, 4G). Similarly, we could observe a striking increase in C-C motif chemokine ligand (CCL) signaling between immune populations in the skin (Fig4E). Of note, we could observe that while expression of the chemokine CCL5 could be found in both skin and circulating T cells populations, expression of the chemokine receptor CCR4 was almost exclusively found among skin T cell types (Fig4F). While the mechanisms driving immune cell infiltration to the skin in BD has not been previously clarified, CCR4 expression has been reported to be a major contributor to skin-homing in the context of other autoimmune diseases and infection^39^. As such, these results suggest that the skin manifestations of BD may involve substantially different immune cell response compared to circulation. Differences in cytokine signaling and cell migration may be particularly important causal drivers of this variation.

**Figure 4.**
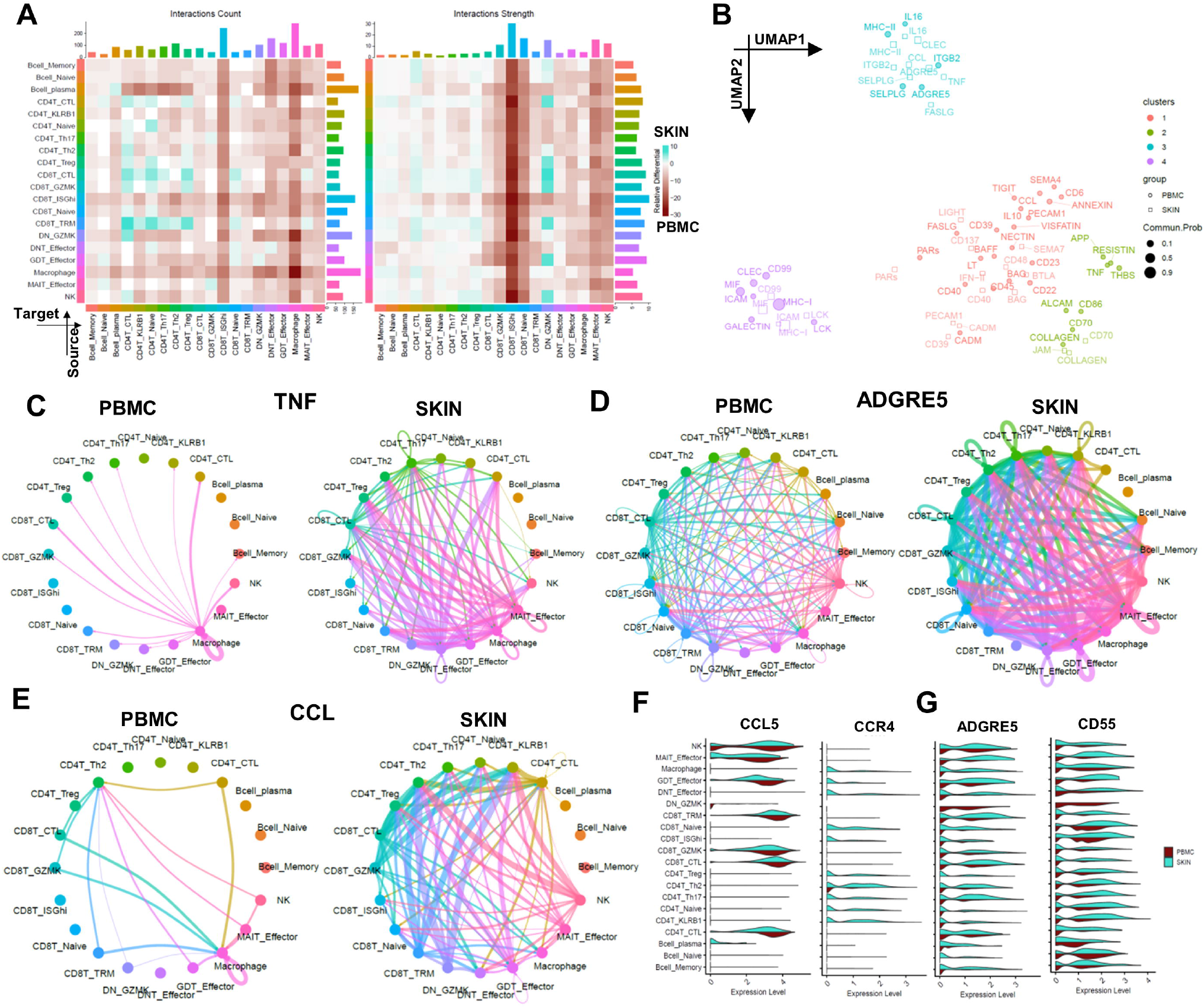
Differential communication characteristics between skin and peripheral immune cells in Behcet’s Disease. A) Heatmap of the interaction matrix between each communication target and source scored based on the total number of distinct interactions found between each pair (left), or the weighted overall strength of all interactions in each pair (right). Garnet grids correspond to interactions between the cell type pair being more prominent among the circulating populations. B) UMAP visualization of the different types of communication interactions found to be significant in at least one tissue, arranged based on functional pattern similarity in this context. C) Circle plot showing the landscape of TNF communication between distinct cell types in circulation (left) and in skin (right). Interaction lines are colored according to the cell type source, while the width of the line corresponds to interaction strength. In the context of TNF, macrophages appear to be the driving source of TNF communication within circulating populations. However, multiple populations of CD4+ and CD8+ T cells are found to also play a role in initiating TNF communication in skin, leading to a heavily interconnected and overlapping communication pattern wherein skin TNF may be anticipated to also be derived from T cell sources. D) Circle plot of the landscape of ADGRE5 communication. ADGRE5 (encoding CD97) engages with CD55 as part of a receptor-ligand interaction between surface molecules. While nearly all cell types are projected to engage in this communication in both skin and circulation, these interactions are much more robust among skin populations. E) Circle plot of the landscape of CCL signaling demonstrates increases in both complexity and interaction intensity in skin populations compared to circulation. F-G) Violin plot of the expression of the CCL5-CCR4 (F) and ADGRE5-CD55 (G) receptor-ligand pairs demonstrates this discordance in communication, as the CCR4 receptor and ADGRE5 ligand are strongly expressed among skin T cell populations but rare in circulating populations.

### Skin-infiltrating and circulating CD8+T share clonal origin and progress to acquire tissue-residential characteristics

While the results above indicate that skin-infiltrating and circulating populations both mount an active immune response in BD patients, matching cell types in the two tissues were also be found to display differential gene expression and cellular crosstalk characteristics. As such, the precise connection between these matching cell types is unclear. To resolve this question, we then utilized TCR repertoire information obtained from concurrent scTCRseq of the same cells to trace the clonal origins of skin-infiltrating T cells. Following maturation in the thymus, the vast majority of mature αβT cells will enter into circulation as naïve cells, with almost every cell displaying a unique TCR sequence corresponding to their antigen specificity. Per common understanding, these naïve αβT cells cannot alter their TCR sequence following thymal egress. However, they may proliferate into a large pool of cells following encounter and activation by their cognate antigens in a process of clonal expansion. Clonally expanded T cells may then distribute across tissues and/or migrate to sites of inflammation to carry out their effector functions. Importantly, during this expansion process, cells of a single clonal lineage may adopt distinct phenotypes as a result of exposure to environmental stimuli (such as cytokines, metabolites, etc.), as well as asymmetric cell division (ACD, wherein a proliferating cell divides into two daughter cells with differing phenotypes). As a result, examination of TCR sequences reads out a genetic record tying all T cells of a single clonal lineage.

Within the CD8+T cell populations we observed in BD patients, we found that the naïve and ISGhi populations in both skin and circulation were composed exclusively of rare, low-frequency clonotypes, indicating that these populations were unlikely to have undergone clonal expansion. However, circulating CTL and Trm populations were dominated by highly expanded clonotypes (Fig5A). These latter two populations were also expanded in the skin, albeit to a lesser extent. Concurrent examination of the transcriptomes of these cells also indicate that GZMK+, CTL, and Trm populations feature elevated cell cycle activity, consistent with expectations that these cells have undergone clonal expansion (FigS7).

**Figure 5.**
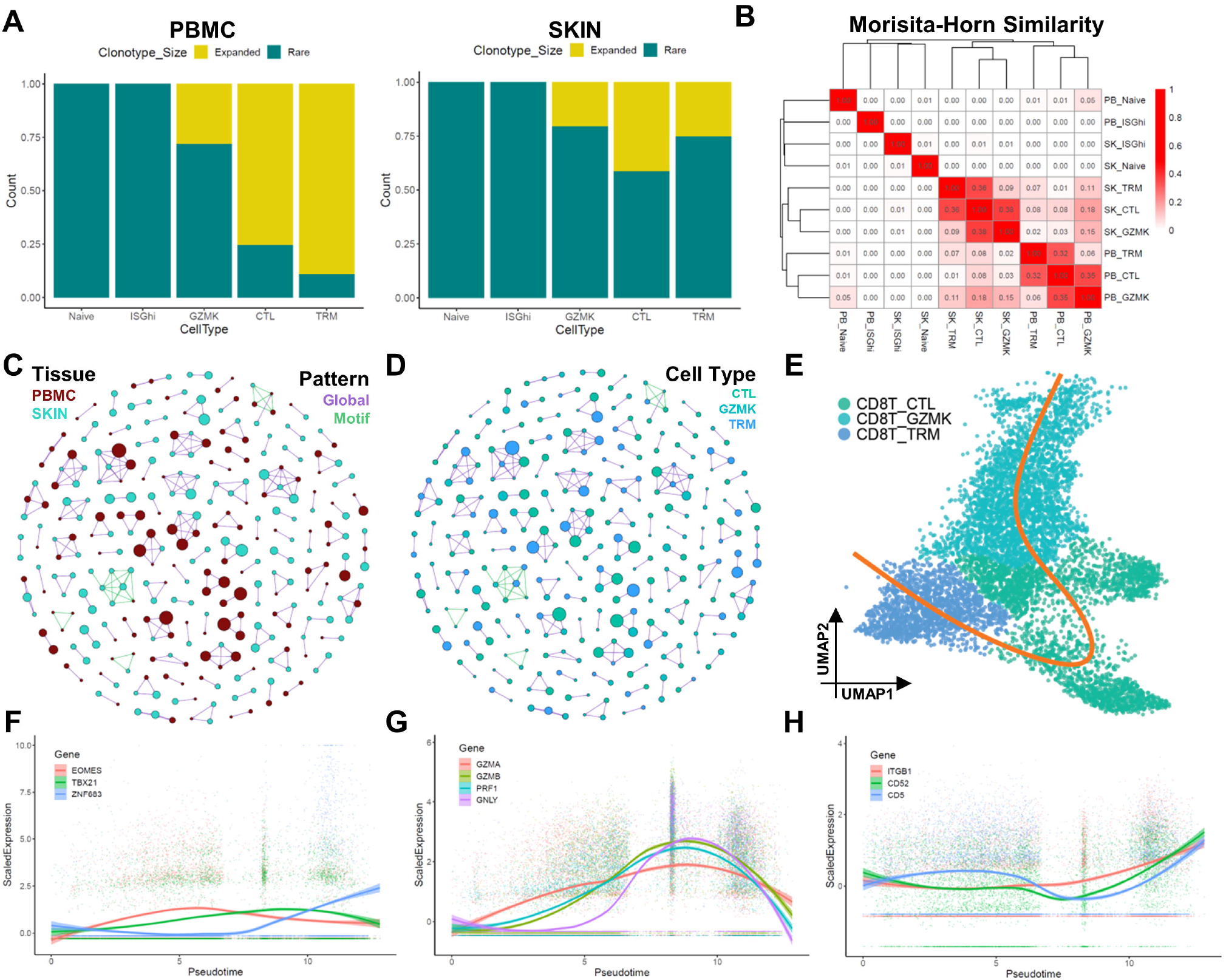
Combined trajectory and repertoire analysis of CD8+T cells identifies clonal sharing between local and circulating memory cells. A) Clonal homeostasis analysis of the five CD8+T cell phenotypes found in circulation (left) and skin (right). In this analysis, clonotypes were considered to be expanded if at least 4 cells with the same TCRβ CDR3 amino acid sequences were found within the same cell type. B) Morista-Horn similarity matrix of TCRβ CDR3 amino acid sequences across skin and circulating CD8+T cell populations, clustered according to similarity. C) Network analysis of TCRβ CDR3 amino acid sequences clustered via paratope hotspots using GLIPH2. Each node corresponds to a single clonotype, with edges connecting clustered clonotypes. Edges are colored by clonotype pattern (global sequence versus motif), while nodes are colored according to tissue origin. Larger nodes correspond to clonotypes of higher frequencies. D) TCRβ network in (C) with nodes colored according to cell type of each node. E) Inferred trajectory line (orange) overlaid in UMAP space of the three CD8+T cell populations showing clonal overlap. The trajectory begins in the CD8+ GZMK population, passes through the CTL population, and terminates in the Trm population. F-H) Gene expression curves of key molecules over the pseudotime trajectory inferred in (E). Three transcription factors show phased changes in expression over the course of the trajectory, culminating in the emergence of ZNF683 in Trm cells (F). Expression of four molecules associated with CD8+T cell killing increase during transition towards CTLs and decrease following the adoption of Trm phenotype (G). Three molecules found to positively increase during the course of CTL to Trm transition (H).

To understand the lineage relation between these differing CD8+T phenotypes, we next assessed the similarity between the disparate populations in skin and circulation. Clustering based on clonotype similarity revealed that circulating and skin-infiltrating GZMK+ and CTL populations featured a large degree of overlap, while naïve and ISGhi populations were composed of distinct repertoire pools (Fig5B). Intriguingly, we also found significant similarity between circulating GZMK+ cells and skin-infiltrating GZMK+, CTL, and Trm populations, and this similarity was found to be supported by substantial numbers of overlapping clones (FigS7, TableS3). Furthermore, clustering of TCRβ sequences across the six cell types with notable clonal expansion revealed that a notable portion of global patterns and sequence motifs could be found to span clonotypes observed in skin and circulation (Fig5C-D). Collectively, these results suggest that memory and effector CD8+T clones in BD patients may disseminate to both skin and circulation, and demonstrate that single clonotypes may efficiently migrate between the two sites. As such, we can infer that differences in transcriptome and communication characteristics do not preclude the same antigenic drivers from influencing BD pathogenesis in both skin and circulation.

To further explore this indication, we then performed trajectory analyses on the CD8+T cell populations showing significant clonal overlap. Interestingly, a single differentiation path was inferred to be the most likely model, wherein GZMK+ memory cells may differentiate into CTLs before then acquiring tissue-resident properties (Fig5E). Consistent with this model, this progression trajectory involves a change in the expression of key transcription factors, with *EOMES* expression decreasing past the GZMK+ memory stage, and *ZNF683* expression being dominant in Trm cells (Fig5F). In contrast to CTL effectors, Trm cells also display lowered expression of cytotoxic effector molecules, consistent with their memory phenotype (Fig5G). Furthermore, the terminal Trm population is enriched for *ITGB1*, an integrin expressed late on during immune response associated with mediating tissue residence, and also show enhanced expression of the *CD5* molecule required for regulating TCR affinity (Fig5H). Taken together, these results indicate that a single pool of memory CD8+T cells mediating immune response in the skin and circulation may diversify to adopt distinctive tissue-specific phenotypes over time. Since these tissue-resident cells may subsequently accelerate regional response upon future exposure to antigen, we hypothesize that this local conversion may contribute to recurrence of inflammation in BD patients, with one round of inflammation laying the groundwork for subsequent recurrence in the future by establishing local residential populations.

### Skin CD4+T cells include a population of expanded Tregs enriched for IL-32 expression

CD4+ T cells are known to produce substantial amounts of cytokines and engage in intracellular communication to maintain inflammatory microenvironments. In order to investigate the factors contributing to CD8+T cells adopting tissue residence in the skin during BD pathogenesis, we next considered the possibility that differentiated CD4+T cell types may play an important role. In particular, we considered the possibility that the expanded population of CD4+T Tregs observed in the skin may play a role, since Tregs are known to influence CD8+T cell behavior^40^. Trajectory analysis of the five CD4+T cell populations identified yielded three distinct paths jointly emerging from naïve CD4+T cells, wherein a portion of cells undergoing an early branching event to progress into CD4+T Tregs, while the remainder branch during a memory-like KLRB1+ stage into Th2 and Th17 cells (Fig6A, FigS8). Closer inspection of the naïve CD4+T cell to Treg path demonstrated that skin Tregs occupied the end stage of this differentiation trajectory, while naïve cells could be more frequently observed in at the start in peripheral circulation (Fig6B-C). These results suggest that the increase in skin Tregs in BD patients may be driven by mechanisms distinct from those regulating Th2 and Th17 populations.

**Figure 6.**
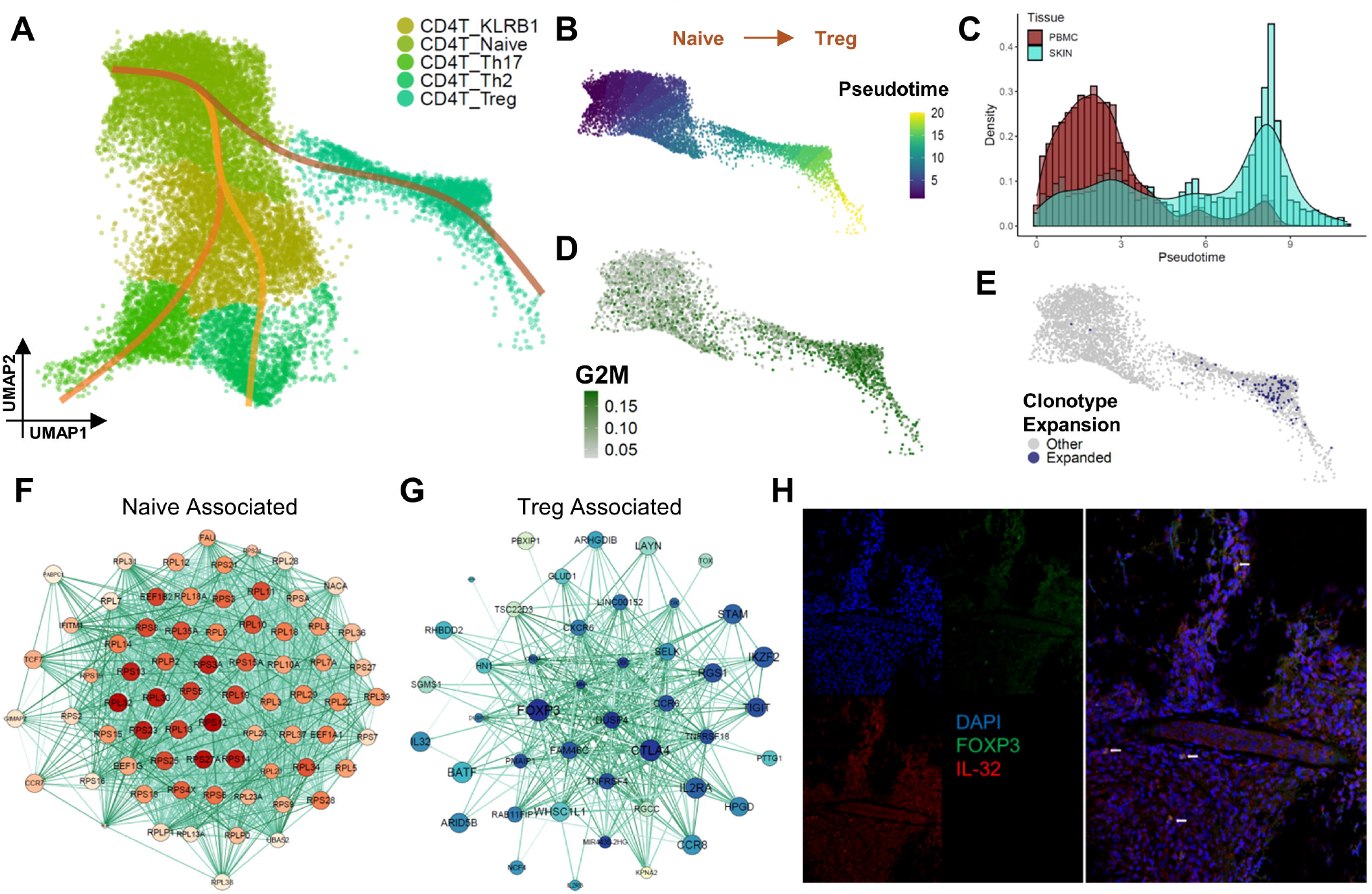
Combined trajectory and repertoire analysis of CD4+T cells identifies clonal expansion of skin Tregs enriched for IL32. A) Trajectory curves inferred among five CD4+T cell populations, with each curve representing a distinct differentiation lineage. B) UMAP visualization of the subset of cells participating in the differentiation trajectory from naïve CD4+T to Tregs, colored according to their assigned pseudotime within this trajectory. C) Histogram showing sample origin of each cell over the pseudotime trajectory marked in (B). D) Cell cycle scoring using G2-M phase genes over the course of the differentiation trajectory in (B) shows notable enrichment of these markers in Tregs. E) Clonotype expansion scoring of the cells in (B). All cells belonging to clonotypes with at least 4 cells within the trajectory were considered to be expanded in this context. F) Correlation network analysis of the top genes showing decreases in expression over the course of naïve to Treg differentiation. Larger nodes correspond to genes with higher mean expression. Darker nodes mark higher network centrality. G) Correlation network analysis as in (F), but of genes showing increases in expression over the course of naïve to Treg differentiation. H) Immunofluorescence staining of FOXP3 and IL32 expression in a skin lesion biopsy of a patient with BD (n = 5). While some background IL32 is apparent, high expression of IL32 notably localizes to cells with positive FOXP3 nuclear staining.

Interestingly, cell cycle scoring along the Treg differentiation trajectory further demonstrated that while naïve T cells were largely quiescent, the Treg population also displayed signs of proliferation (Fig6D). To clarify this point, we then performed TCR repertoire analysis. Intriguingly, these skin Tregs included a portion of cells of common clonal origin and matching TCRβ sequences, a phenomena not seen in circulating Tregs (FigS9, FigS6E), indicating that these cells are highly likely to have truly proliferated. Clonotype similarity analysis further demonstrated that skin Treg clonotypes show a slight overlap with skin naïve cells and circulating Tregs, but no overlap with circulating naïve cells (FigS9). Unfortunately, the repertoire similarity and overlap levels among CD4+ T cell populations were notably lower than those observed in CD8+T cells, and expanded clone sizes were also smaller, making it difficult to draw further inferences regarding their clonal origin. Nonetheless, these results raise the possibility that the expansion in skin Tregs observed in BD patients may be influenced by local proliferation.

To further investigate the origin of these expanded Treg cells, we then carefully examined their transcriptome signatures. Network analysis of the factors most strongly positively associated with progression towards the Treg phenotype demonstrated that naïve cells were primarily enriched for ribosome components, together with the surface receptor CCR7 and transcription factor TCF1, and that expression of these factors declined over time (Fig6F). In contrast, progression to the Treg phenotype included a change in several transcription factors, including BATF, IKZF2, and FOXP3 (Fig6G). Interestingly, we also observed prominent expression of the cytokine IL32 to be associated with Treg progression, and Treg expression of this cytokine was the highest compared to other cell types (FigS10). At the same time, these Tregs did not appear to express significant levels of inhibitory factors, such as IL10. To validate the expression of IL32 in skin Tregs, we then performed immunofluorescent staining of skin biopsy samples from five patients with BD. Consistent with the sequencing results, we observed strong IL32 signal in FOXP3+ skin Tregs (Fig6H). Collectively, these results suggest that skin Tregs in BD represent a distinctive cell population with unique functional characteristics.

### IL32 does not enhance regulatory T cell function but increases CD97 signaling

Tregs have been demonstrated to exert their regulatory function through a number of different molecular mechanisms, including secretion of cytokines such as IL-10 and TGFβ, increasing environmental adenosine levels via CD39 and CD73, and suppressing antigen-presenting cell function via surface ligands such as CTLA-4 and LAG3^41^. Tregs are also widely appreciated to exert their suppressive functions in both circulation and at tissue sites, with tissue Tregs being understood to be primed towards homeostatic responses under physiological conditions^42^. However, the role of IL-32 secretion in Tregs has not been clearly defined. In order to further explore the functional significance of the production of this cytokine, we first examined its possible impact on T cell behavior. Initial differentiation experiments demonstrated that IL32 did not have a substantive effect on the proportion of Tregs induced from PBMCs isolated from healthy donors (Fig7A). Furthermore, in vitro addition of IL32 to cultures of PBMCs isolated from healthy donors did not appreciably impact the proliferative ability of either conventional CD4+T or CD8+T cells in response to canonical TCR activation (Fig7B). At the same time, IL32 also failed to show a significant impact the expression of key pro-inflammatory cytokines such as TNF or IFNγ (Fig7C). Taken together, these results indicate that IL32 derived from Tregs is unlikely to have an inhibitory effect on T cell function.

**Figure 7.**
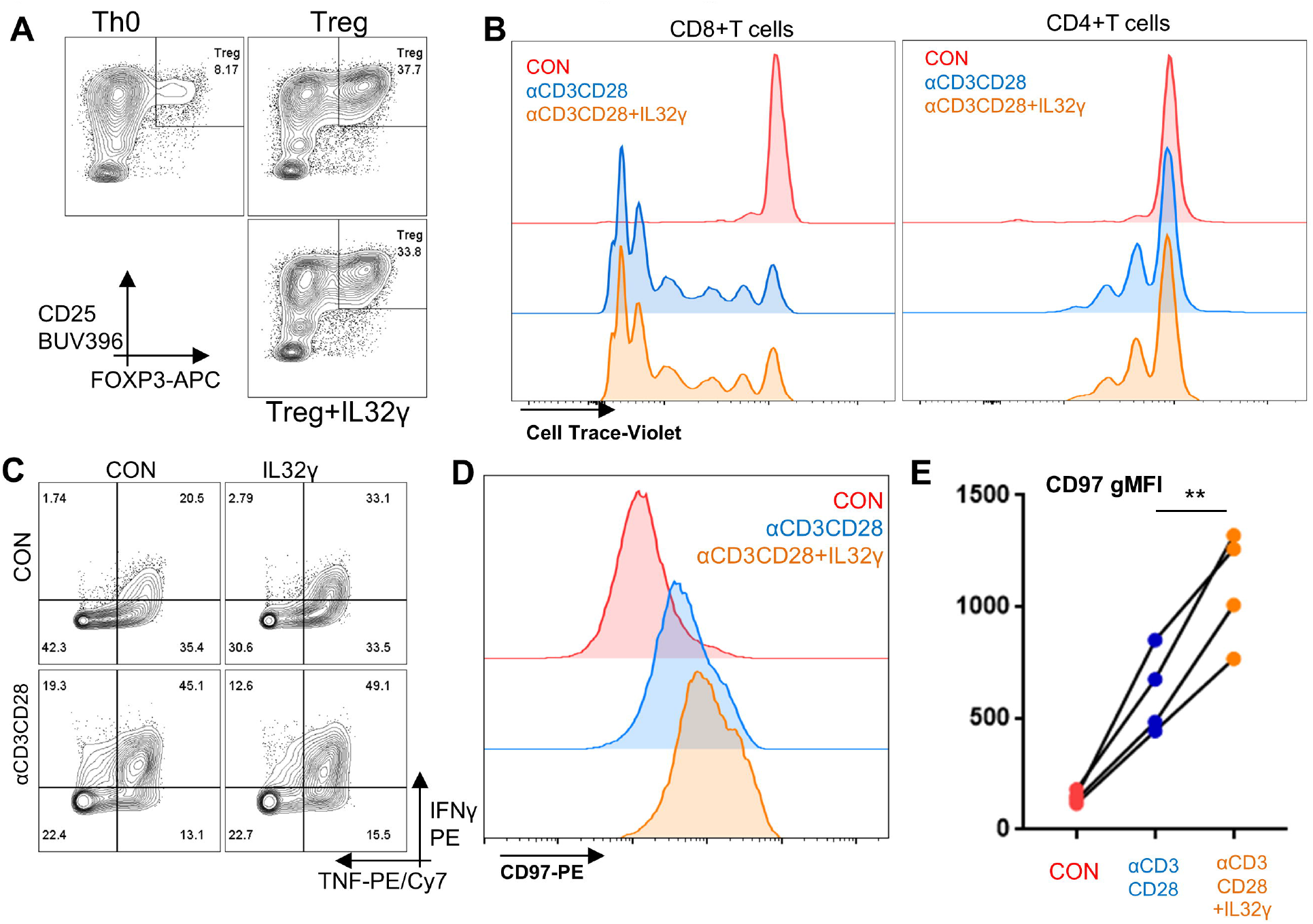
IL32γ does not suppress CD8+T cell proliferation or cytokine production but increases CD97 signaling. A) Differentiation of CD4+T cells derived from human PBMCs for three days in the presence or absence of IL32γ shows no significant change in the percentage of FOXP3+CD25hi Tregs (n = 4). B) Cell proliferation histograms of conventional CD8+T (left) and CD4+T (right) cells labeled with membrane dye and activated by TCR stimulation via αCD3CD28 for 6 days (n = 8). No significant distribution change was elicited by IL32γ. C) Intracellular staining of cytokine production in CD8+T cells activated by TCR stimulation via αCD3CD28 for 3 days (n = 8). IL32γ could not restrain either baseline or stimulation-induced cytokine production. D) Histogram of CD97 expression in CD8+T cells 6 days post-activation shows prominent rightward shift in CD97 expression resulting from IL32γ treatment (n = 4). E) Statistical summary of the geometric mean fluorescence intensity of CD8+T cells from (D). p < 0.01.

Since IL32 did not show a suppressive impact on T cell cytokine production, we then examined the possibility that it may impact the intracellular communication networks we had observed to vary between skin and peripheral blood. Expression of the chemokine receptor CCR4 and the skin-homing marker CLA were not notably effected by IL-32 stimulation (data not shown). However, we found that prolonged stimulation with IL32 highly increased T cell surface expression of CD97 (Fig7D). Of note, this striking increase came on top of the increase caused by canonical TCR activation (Fig7E). This increase coincided with a significant increase in CD97-CD55 mediated communication between skin immune populations relative to peripheral circulation seen earlier. Collectively, these results thus indicate that not only does IL32 not have a direct suppressive effect on T cell responses; it may also contribute to T cell infiltration and tissue invasion through CD97 signaling, and accelerate BD pathogenesis.

## Discussion

Defects in both the relative proportion of Tregs, and individual suppressive function of Tregs to restrain CD4+ and CD8+ T cell-mediated immune responses, have been widely reported to occur in a number of autoimmune diseases, including various types of vasculitis^43,44^. A number of previous reports have sought to address the changes in relative proportion of circulating Tregs in BD patients, generally showing of decreases^45,46,47^ in Treg percentages in active BD patients relative to healthy control. Treg percentages in BD patients have also been shown to change substantially in response to treatment^48,49^. However, whether the decrease in Tregs also extends to sites of BD tissue manifestation has not been previously demonstrated, and the functional capabilities of Tregs in the context of BD have not been clearly examined. From our results, we observe that the percentage Tregs is in fact quite high in the skin of patients with BD when compared to circulation. Furthermore, we observed that these skin Tregs showed signs of clonal expansion, and expressed high levels of the cytokine IL32.

High Treg expression of IL32 has been previously reported to be positively associated with inflammation in the context SARS-Cov-2 infection^50^, a condition wherein Tregs have been shown to display dramatic perturbations^51^, and has been suggested to permit Tregs to carry out pro-inflammatory functions. This latter presumption has been supported by abundant evidence in innate immune cells that IL32 can induce the differentiation of monocytes into mature macrophages^52^, while also stimulating the expression of costimulatory molecules and cytokines in macrophage and dendritic cell populations^53,54^. These effects have also been showed to indirectly influence T cell behavior, particularly in terms of CD4+T cell differentiation^55^. From our results, we found that IL32 also holds an additional capability in increasing the expression of CD97 on activated CD8+T cells, and CD97 signaling was elevated in skin-infiltrating CD8+T cell populations. These indications thus suggest that the high proportions of IL32-expressing Tregs are likely to be dysfunctional, and instead act as positive regulators of tissue damage in the skin of BD patients over time. Further validation work into this possibility may shed more light on the full importance of these Tregs and other CD4+T cell types to both vascular and dermal pathogenesis in BD.

Although mutations in HLA-B*51 have long been appreciated to be strongly associated with BD, the function and trajectories followed by the CD8+T cells that survey and recognize peptides displayed on class I MHC molecules has not explored in detail in this context. Through our integration of scTCRseq and scRNAseq data, we observed that CD8+T cells largely fell into three classes of effector (GZMK+, CTL, and Trm) phenotypes in both circulation and skin manifestations of BD patients. These phenotypes are intimately linked by TCR clonal identity, suggesting that the same T cell clones are capable of mediating effector function in both tissues. We further identify through trajectory analysis a differentiation trajectory followed by these clones, wherein cytotoxic effector cells may adopt tissue-resident characteristics during active disease, and consequently establish themselves into a local memory population to more rapidly respond to future stimuli. While the contribution of Trm cells in the context of BD has been hitherto underappreciated, Trm have been found to mediate long-term antigen memory^56^ and the recurrence of inflammation in multiple skin diseases^57^. Taken together, our data can thus be synthesized into a model for BD recurrence (Fig8) wherein IL32 derived from Tregs may boost the adoption of resident characteristics by tissue-infiltrating CD8+T cells during an initial inflammatory episode. Prolonged residence of these CD8+T cells may then contribute to repeated recurrence of skin lesions in response to future stimuli. Further investigations can clarify the full contribution of these Trm cells and their dependence on Treg function in relation BD recurrence, and also address whether the same clones will repeatedly function as the protagonists across multiple rounds of active disease.

**Figure 8.**
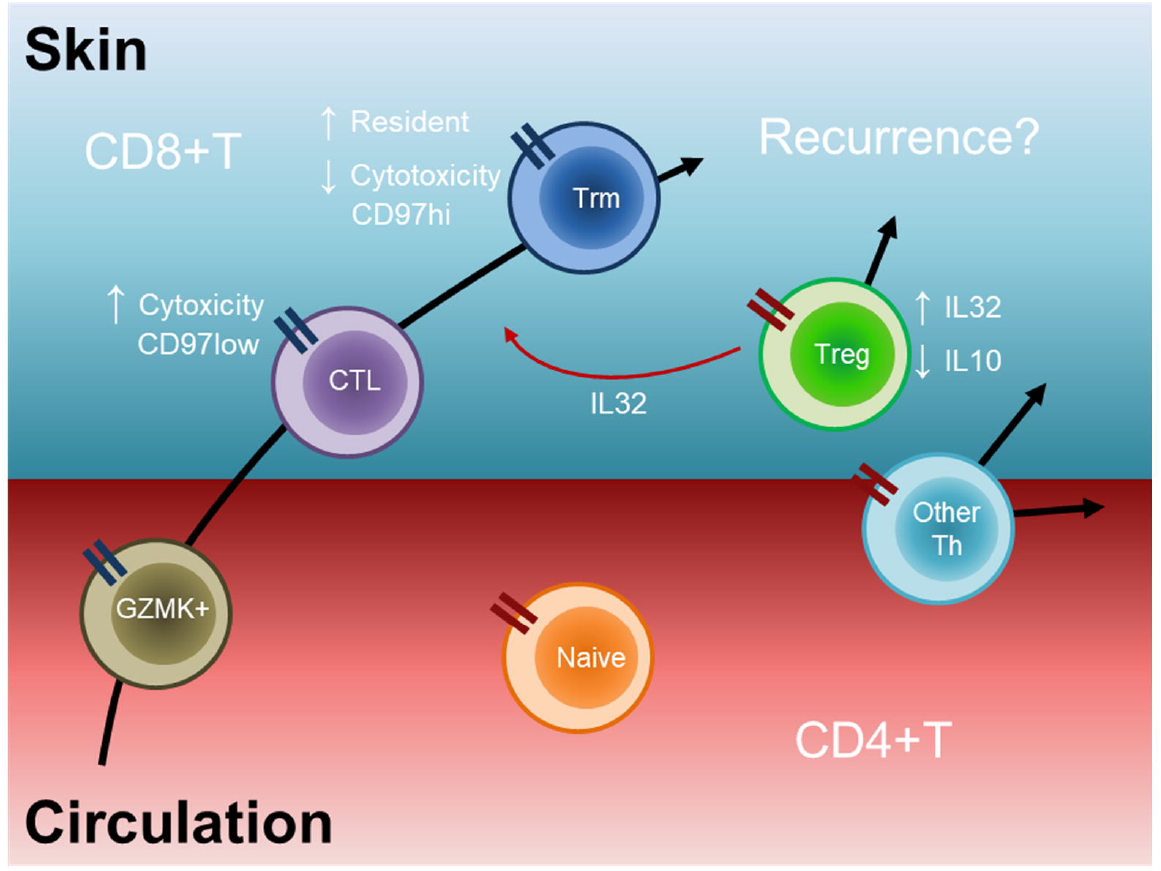
Model of T cell dynamics driving recurrence of BD. By synthesizing the trajectories followed by CD4+T and CD8+T cells in BD, we propose a model wherein CD8+T cells progress towards the acquisition of resident memory properties during an inflammatory episode. This progression coincides with a progression of naïve CD4+T cells towards an aberrant Treg phenotype overexpressing IL32. IL32 from Tregs may further support the acquisition of Trm phenotype via CD97 expression. These Treg and Trm populations may persist through resolution of the inflammatory episode, and contribute to rapid responses in future episodes of skin inflammation.

In addition, identification of the antigen specificity of these CD8+Trm may help to significantly advance our understanding of the cell targets of CD8+T cells in the skin and other affected sites. While our study identifies a number of expanded Trm clones that may be autoreactive, we were unable to determine their antigen sensitivity. One report has found that a portion of BD patients possess circulating CD8+T cells that are responsive to a peptide in MICA as presented by HLA-B*51^58^, while other autoantigens from IRBP might also be sensed^59^. Longitudinal tracking in BD patients with classified antigen specificities may serve to weave together these strands into a complete tapestry of CD8+T cell function in BD.

While scRNAseq has immense potential for unraveling cell type heterogeneity and functional variations in complex environments, it should also be noted that current single-cell technologies also contain inherent limitations. For instance, in our current study, we elected to focus on CD45+ mononuclear cells in order to optimize cell capture efficiency, thus excluding stromal cells in the skin, as well as granulocytes, which may also play critical roles in BD pathogenesis^60^. Droplet-based sequencing techniques may also be somewhat limited their ability to capture transcripts with low expression levels^61^. As such, future studies that leverage more sensitive detection techniques and sample other cell types may be able to further improve the census resolution. Furthermore, while our study identified substantial numbers of regulatory and expression differences in immune response between peripheral circulation and skin lesions, it is unclear how many of these differences may also extend to other site of BD tissue manifestation, such as mucosal ulcers, or the nervous system. Intriguingly, IL32 may be a shared element, with a recent report having shown increased IL32 levels in the cerebrospinal fluid of BD patients with neurological symptoms^62^. Further evaluations with longitudinal sampling at other tissue sites may be able to address these and other questions regarding BD pathogenesis.

## Acknowledgements

The authors would like to thank all lab members for their helpful input during discussion of this work. This work was funded in part by support from the National Natural Science Foundation of China (81971546 and 82171787 to L.Z.) and the Chongqing International Institute for Immunology (2020YJC06 to L.Z.).

