## Supplementary Figures for "Single cell Transcriptome and T cell Repertoire Mapping of the Mechanistic Signatures and T cell Trajectories Contributing to Vascular and Dermal Manifestations of Behcet’s Disease"

### **FigureS1–** Single cell RNA QC

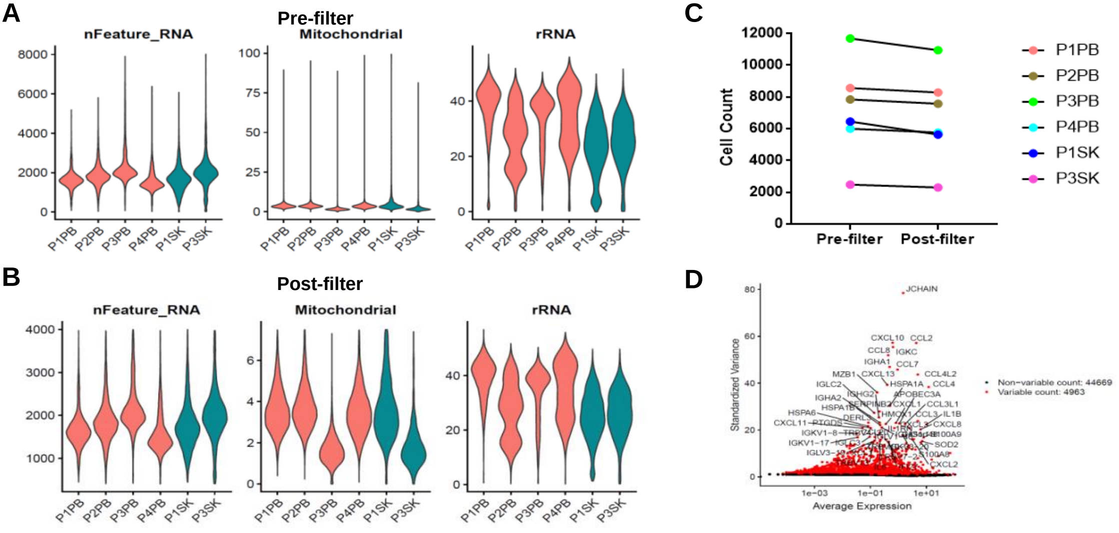

### **FigureS2–** Enrichment of IgA-producing memory B cells in BD

**A**

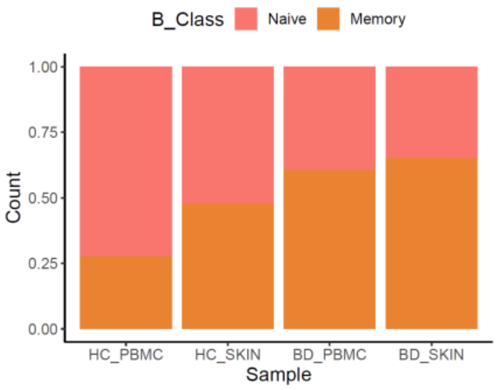


**B**

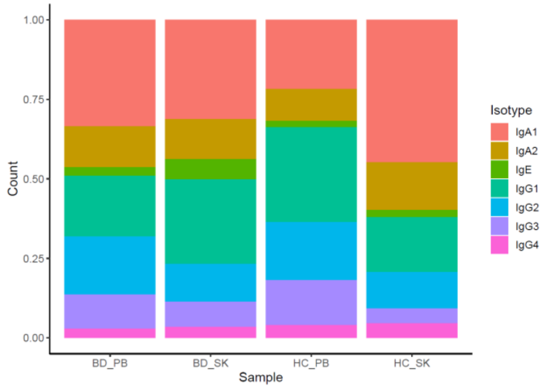


**C**

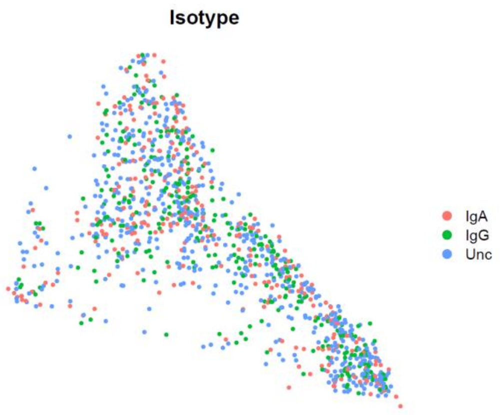

**FigureS3–** Average expression of genetic association markers in skin and peripheral immune cell populations

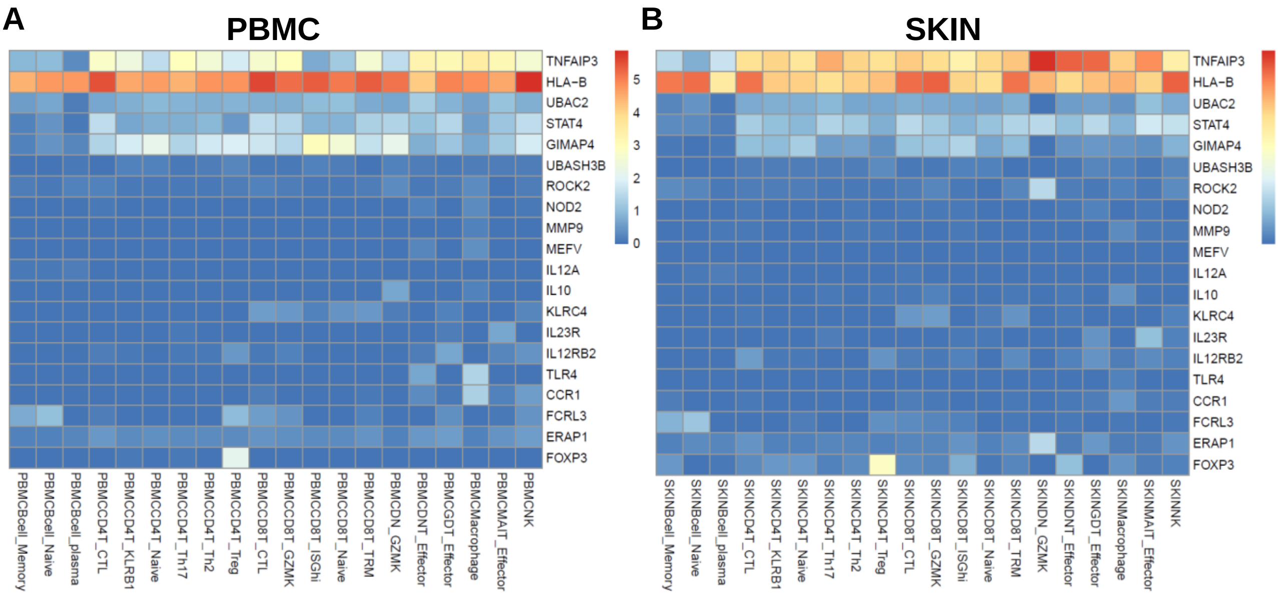

**FigureS4–** DEGs between circulating and skin immune cell populations

A

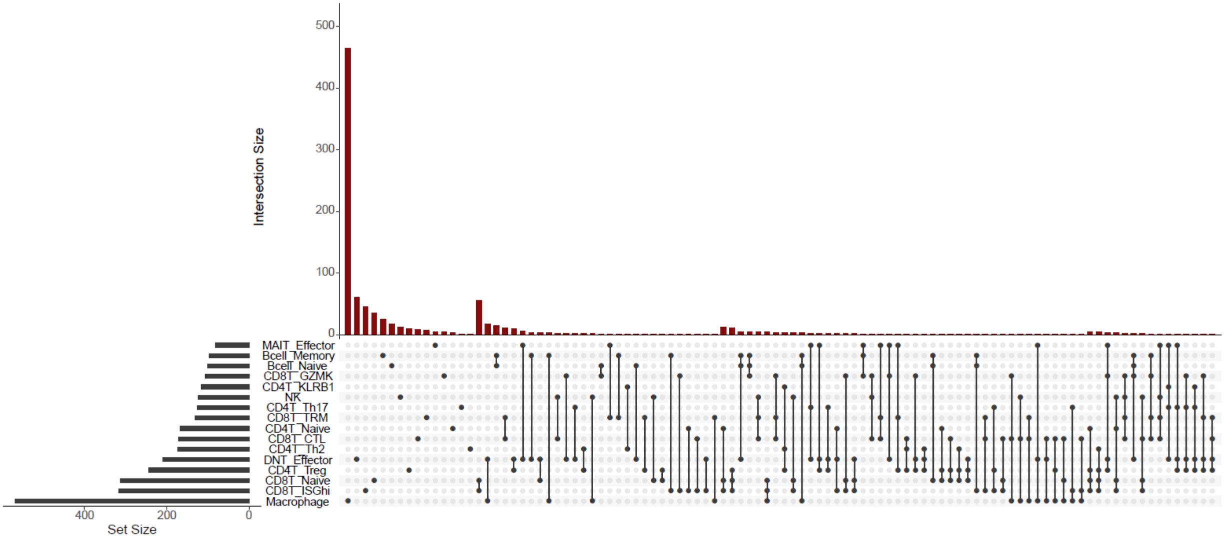


B

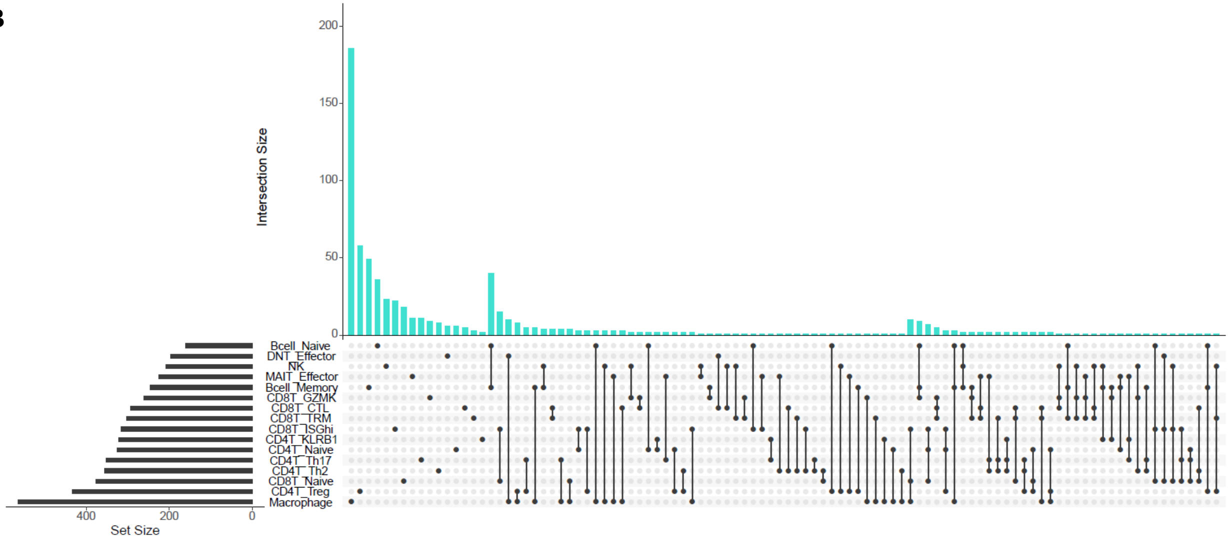

**FigureS5–** Communication analysis identifies significant variance between Circulating and Skin immune populations in Behcet's Disease

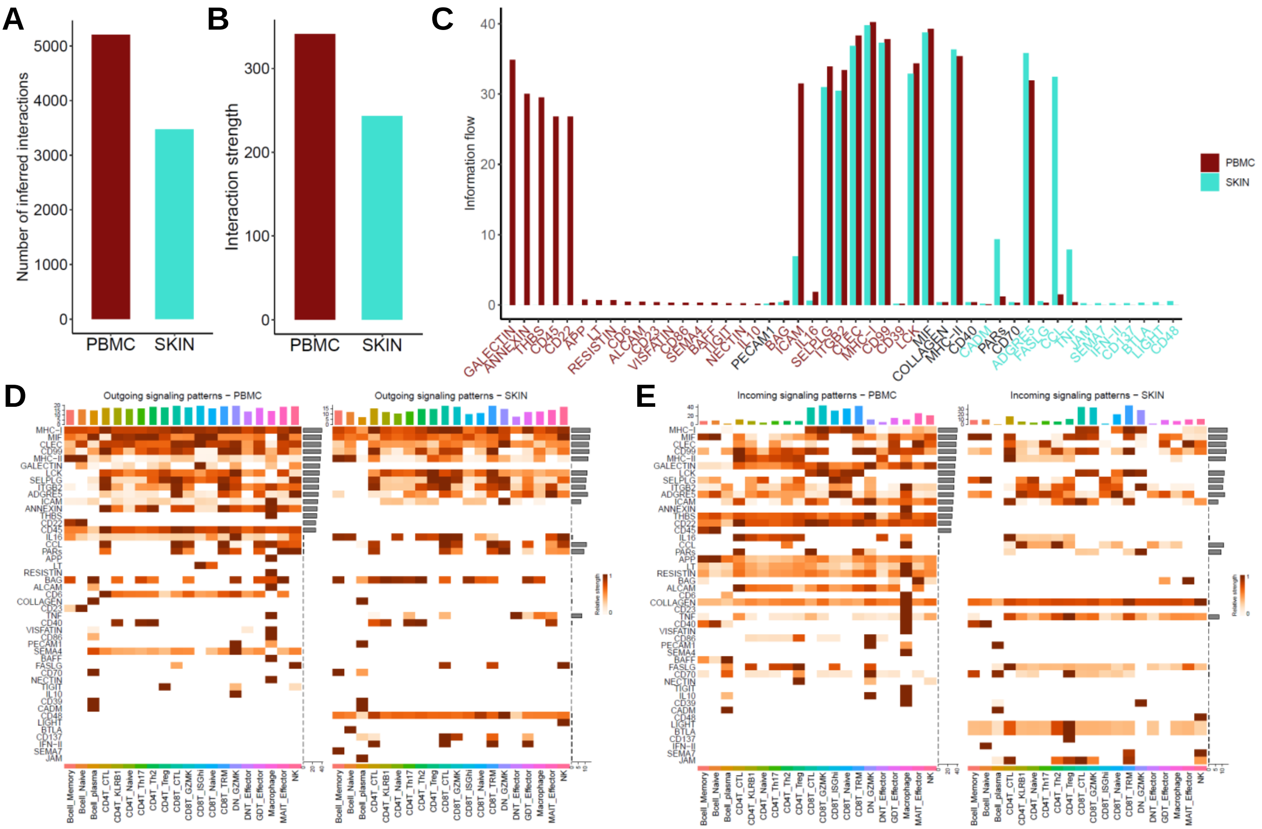

**FigureS6–** Specific enriched R-L pairs in skin and peripheral immune cell populations

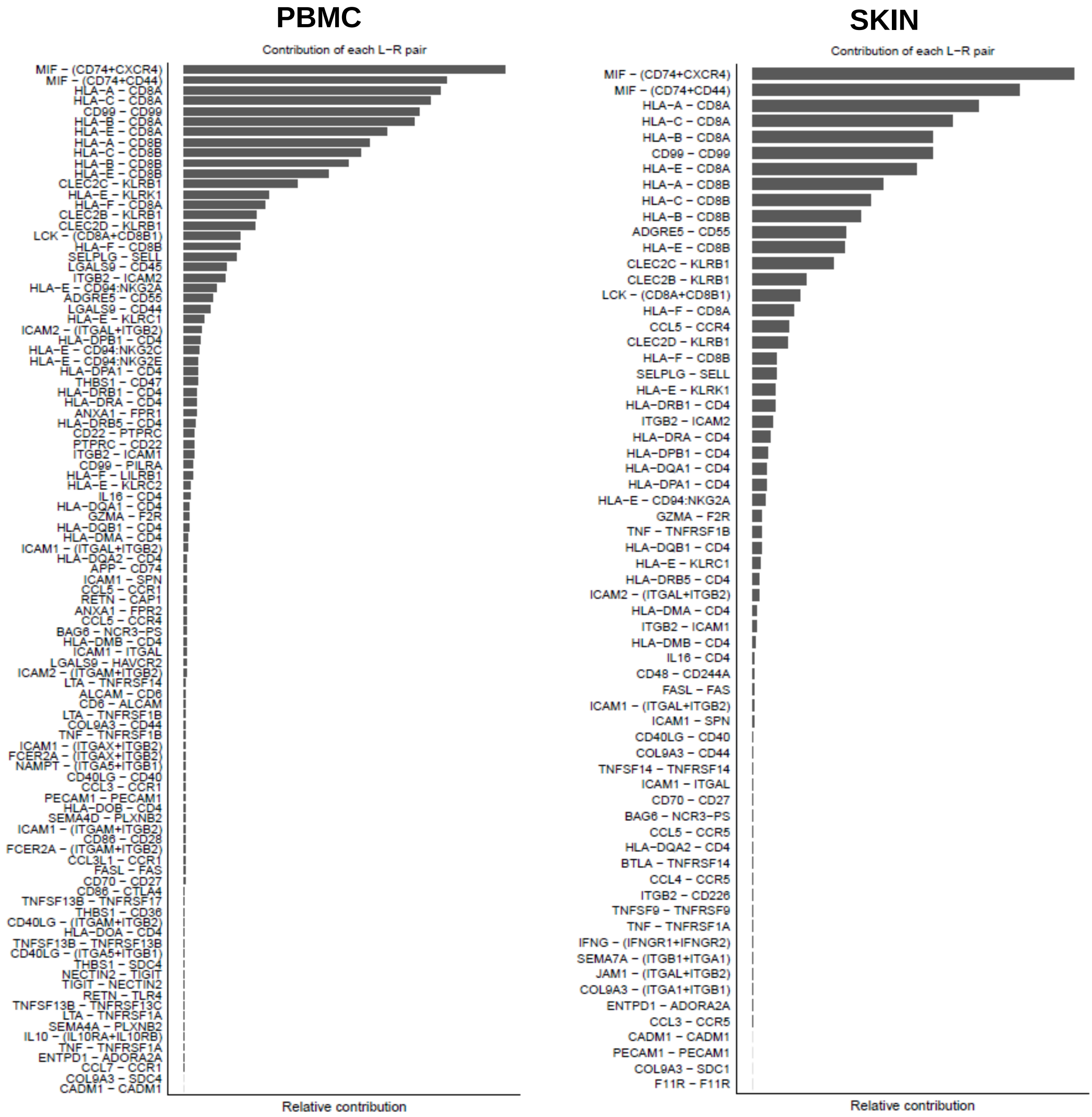

**FigureS7–** TCR repertoire of CD8+T cell populations

A

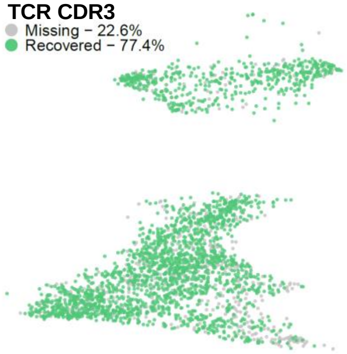


B

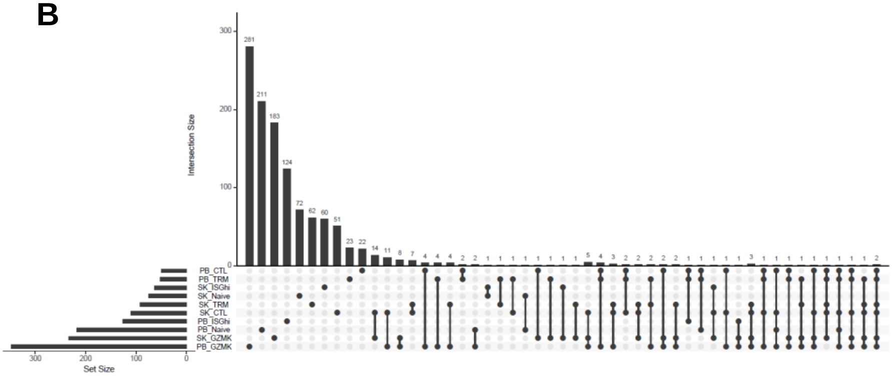

**FigureS8–** Differentiation trajectories of CD4+T cell populations

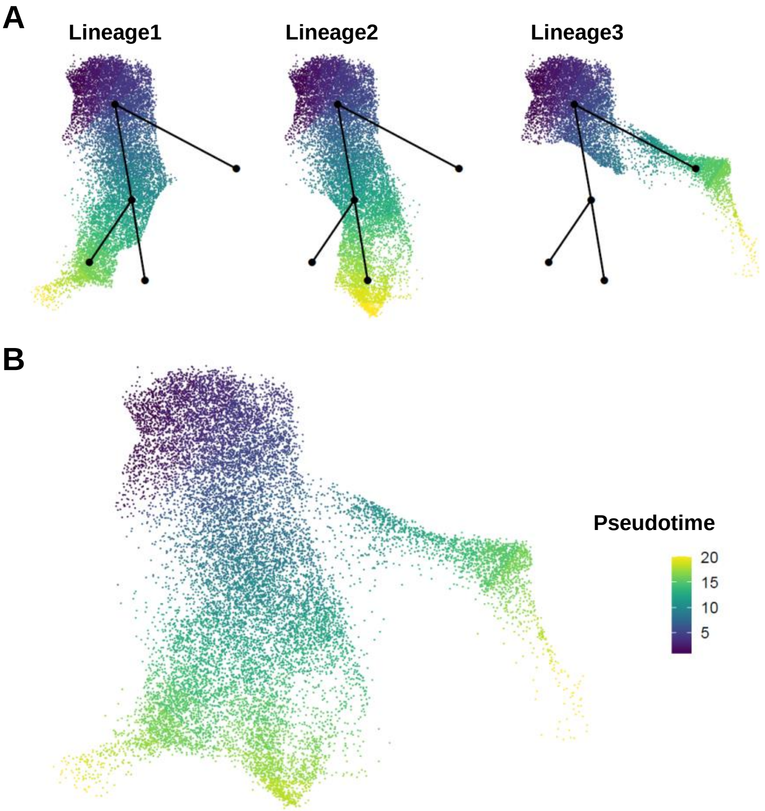

**FigureS9–** TCR repertoire of CD4+T cell populations

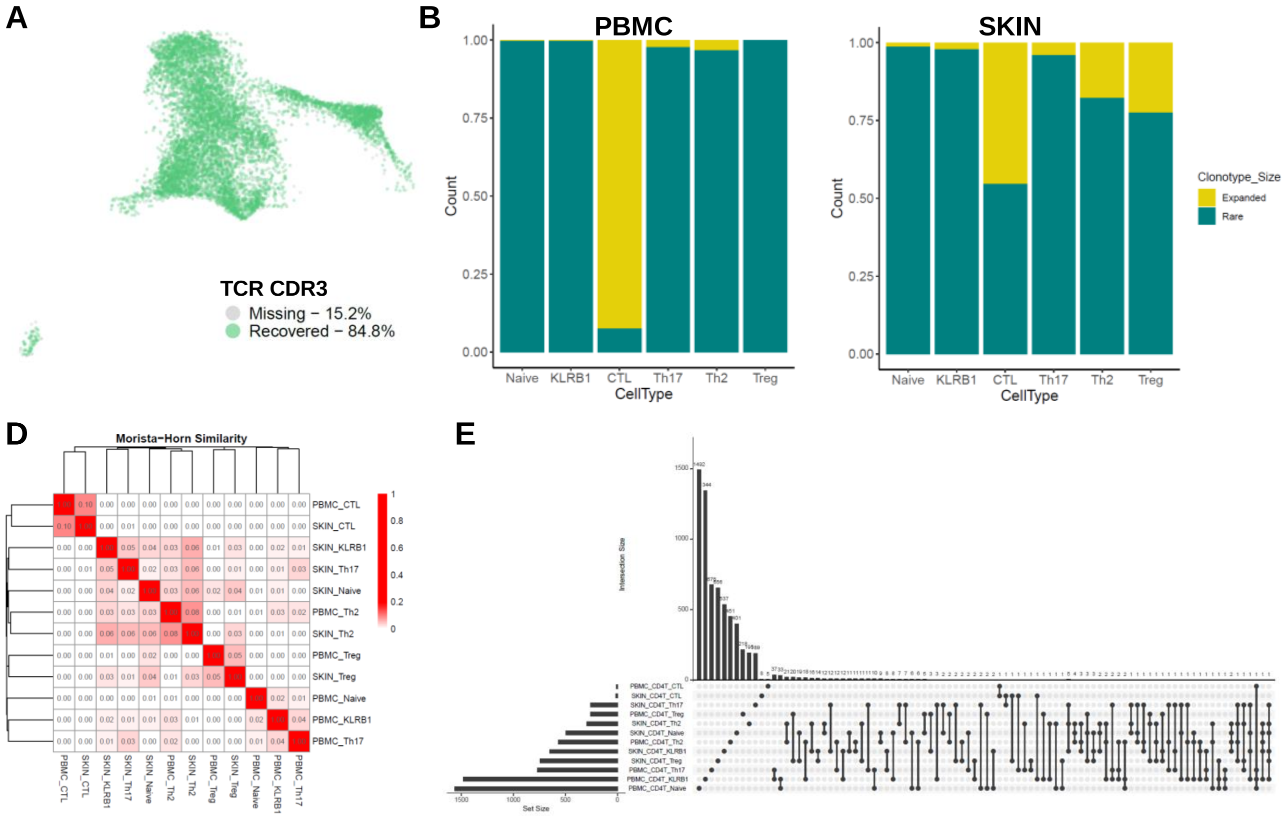

**FigureS10–** Average expression of key cytokines by cell type

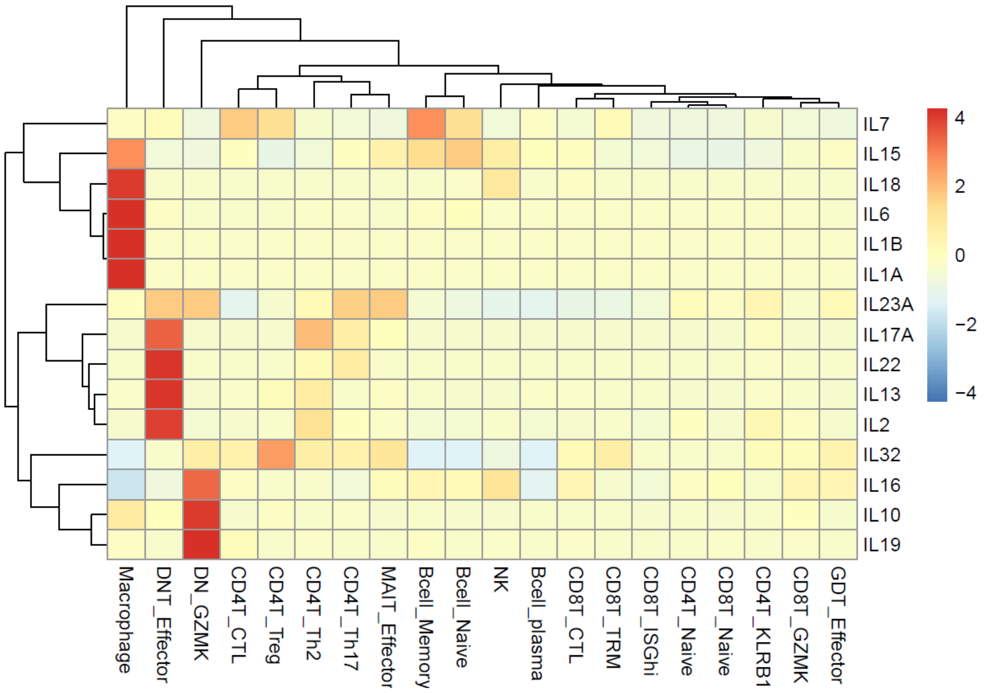
