## Supplementary Figure Legends for "Single cell Transcriptome and T cell Repertoire Mapping of the Mechanistic Signatures and T cell Trajectories Contributing to Vascular and Dermal Manifestations of Behcet’s Disease"

Figure S1—scRNAseq QC. A-B) Violin plots showing the distribution in each cell of the number of genes detected (left), percentages of reads corresponding to mitochondrial RNA (mid), and percentage of reads corresponding to ribosomal RNA (right) before filtering (A) and after filtering based on stringent criteria (B). Cells with >7.5% mt-RNA reads were discarded, as were cells with >4000 or <400 genes detected. Feature detected in fewer than 10 cells in total were also discarded. These quality control steps filtered the dataset down to 39,820 high-quality single cells. C) Line plot showing the impact of filtering on each sample in terms of cell count. Overall quality in each sample was similar. D) Scatterplot of the average expression and expression variance for all features analyzed. The top 10% of variable features returned through vst were retained for subsequent integration.

Figure S2—Enrichment of IgA-producing memory B cells in BD. A) Bar plot of the relative proportion of naïve to memory B cells in healthy PBMC and skin (derived from GSE165816) and our dataset of PBMC and skin from BD patients. Notably, the proportion of memory cells is significantly higher in BD patients in both tissues. Bar plot of the isotype identities memory B cells in each sample. An increase in IgA producing clones could be seen in circulation of BD patients compared to healthy control. C) Visualization in UMAP space of IgA+ versus IgG+ memory B cells in BD patients does not show notable differences in distribution, with both being highly admixed.

Figure S3— Average expression of genetic association markers in skin and peripheral immune cell populations. A) Heatmap of the normalized expression of genetic association markers averaged within each cell type in peripheral blood. Genes are arranged in descending order based on hierarchical clustering of overall expression in both skin and peripheral blood. B) Heatmap as in (A) of the normalized expression of genetic association markers averaged within each cell type in skin. Row and column order is consistent in both plots for ease of comparison.

Figure S4— DEGs between circulating and skin immune cell populations. A) Upset intersection plot of the genes showing positive differential expression in a circulating population as compared to the matching population in skin. B) Upset intersection plot of the genes showing positive differential expression in a skin population as compared to the matching population in circulation. While some genes are consistently altered at significant levels across multiple populations, the large majority are unique to a specific population, indicating that each population may be trained in a distinctive fashion by the differential immune microenvironment in the skin.

Figure S5— Communication analysis identifies significant variance between Circulating and Skin immune populations in Behcet’s Disease. A-B) Total number of interactions inferred to occur in circulating populations and their weighted interaction strength is higher than in skin. C) Bar graph of information flow reveals that a number of interactions are not clearly captured in skin, including those involving galectins, annexins, and CD22. However, we also observe substantially stronger interactions in ADGRE5, CCL, and TNF in the skin among interactions significantly detected in both tissues. Interaction name is colored according to direction of significance (p < 0.05), with names in black being not significant. D-E) Heatmap visualization of the specific interaction patterns of each cell population in circulation (left) and in skin (right). Outgoing interactions are marked in (D), while incoming interactions are shown in (E).

Figure S6— Specific enriched R-L pairs in skin and peripheral immune cell populations. Relative communication intensity of each contributing receptor-ligand pair in circulation (left) and skin (right).

Figure S7— TCR repertoire of CD8+T cell populations. A) UMAP visualization of CD8+T cells from a single patient (both skin and circulating). Over 75% of these T cells had a recovered clean TCR sequence, with recovered cells being uniformly distributed. B) Intersection analysis of TCRβ amino acid clonotypes across all five CD8+T cell populations in both skin and circulation.

Figure S8— Differentiation trajectories of CD4+T cell populations. A) Trajectory analysis using slingshot inferred the presence of three distinct differentiation trajectories in CD4+T cells following three lineages. Of note, an individual cell may fall into one or more lineages, and be positioned in slightly different pseudotime states accordingly. As such, each lineage is analyzed separately to prevent confusion, while an averaged plot marking the general progression rate of all three lineages is given in (B). The population of CD4+ CTLs is not included in this analysis, as that population is located at a large distance in UMAP space, and any intermediary states that may be required for progression to CTL fate were not clearly observed.

Figure S9— TCR repertoire of CD4+T cell populations. A) UMAP visualization of CD4+T cells from a single patient (both skin and circulating). Over 80% of these T cells had a recovered clean TCR sequence, with recovered cells being uniformly distributed. B) Clonal homeostasis analysis demonstrates that CTLs are highly expanded in both circulation and skin. Interestingly however, skin Th2 and Tregs are uniquely expanded compared to circulation. A clonotype composed of >4 cells of the same fate is considered to be expanded. D) Morista-Horn similarity across all sixCD8+T cell populations in both skin and circulation. Interestingly, CTLs in skin and circulation share some clonal overlap, but most other populations only share clonotypes at very low levels (especially relative to CD8+T cell populations seen in Figure5). E) Intersection analysis of TCRβ amino acid clonotypes across all six CD4+T cell populations in both skin and circulation.

Figure S10—Average expression of key cytokines by cell type. Heatmap of the averaged expression profiles of the interleukin molecules detected in at least one cell population and clustered in both directions. Notably, Tregs have the highest average IL32 expression of all cell types, while the traditional Treg inhibitory cytokine IL10 is in fact found at high levels in a population of double-negative T cells expressing GZMK, and to a lesser extent in macrophages, but not significantly expressed in Tregs.
